# Shifts in leaf senescence across the Northern Hemisphere in response to seasonal warming

**DOI:** 10.1101/2021.07.23.453498

**Authors:** Lei Chen, Sergio Rossi, Nicholas G. Smith, Jianquan Liu

**Author notes:** Email of corresponding author: Lei Chen, Jianquan Liu.

## Abstract

Shifts in plant phenology under ongoing warming affect global vegetation dynamics and carbon assimilation of the biomes. The response of leaf senescence to climate is crucial for predicting changes in the physiological processes of trees at ecosystem scale. We used long-term ground observations, phenological metrics derived from PhenoCam, and satellite imagery of the Northern Hemisphere to show that the timings of leaf senescence can advance or delay in case of warming occurring at the beginning (before June) or during (after June) the main growing season, respectively. Flux data demonstrated that net photosynthetic carbon assimilation converted from positive to negative at the end of June. These findings suggest that leaf senescence is driven by carbon assimilation and nutrient resorption at different growth stages of leaves. Our results provide new insights into understanding and modelling autumn phenology and carbon cycling under warming scenarios.

## INTRODUCTION

Tree phenology mirrors the timing of budburst, leaf-out, flowering, leaf senescence and other related biological events (Richardson *et al*. 2013; Piao *et al*. 2019). Shifts in tree phenology alter the length of the growing season (Cleland *et al*. 2007; Richardson *et al*. 2018b) and influence the productivity of terrestrial forest ecosystems (Richardson *et al*. 2010; Zhang *et al*. 2020). Tree phenology also drives water-energy balances and trophic interactions (Edwards & Richardson 2004; Peñuelas & Filella 2009; Richardson *et al*. 2013; Thackeray *et al*. 2016). Phenological changes in trees therefore provide a clear, visible signal of, and an important basis for modelling, how global warming influences terrestrial ecosystems (Xia *et al*. 2015; Chuine & Régnière 2017; Zhang *et al*. 2020). There is now considerable evidence that climate warming has altered tree phenology (Piao *et al*. 2019; Chen *et al*. 2020; Menzel *et al*. 2020). For example, advances in the dates of spring leaf-out in response to warming have been consistently observed over recent decades (Wolkovich *et al*. 2012; Fu *et al*. 2015; Chen *et al*. 2018). However, responses of autumn leaf senescence in temperate trees to warming are idiosyncratic. Both advanced and delayed trends in leaf senescence have been reported under warming conditions (Menzel *et al*. 2006; Jeong *et al*. 2011; Gill *et al*. 2015). The ecological mechanisms underlying these contradictory warming responses remain unclear, making it difficult to predict how the effects of global warming on leaf senescence in trees will impact forest ecosystem functioning in the future (Richardson *et al*. 2010; Piao *et al*. 2019; Chen *et al*. 2020; Jeong 2020; Zhang *et al*. 2020).

The final stage through which tree leaves pass before death is accompanied by the degradation of various macromolecules (e.g., chlorophyll and other proteins) (Woo *et al*. 2013). The main function of leaves at this late stage of the season is to remobilize nutrients (e.g., nitrogen and phosphorus) from aging leaves into perennial trunks, twigs and roots for overwintering and to support growth in the following spring (Vergutz *et al*. 2012). Trees have been shown to delay leaf senescence in order to remobilize more nutrients from old leaves (Estiarte & Penuelas 2015). The progress of leaf senescence therefore depends on whether nutrients have been resorbed to their maximum potential extent. However, the timing of leaf senescence is also determined by the maximum amount of assimilated carbon that can be stored (or sink limitation of photosynthesis) early in the growing season (Paul & Foyer 2001). If warming (or other factors, e.g., elevated carbon dioxide and light levels) speeds up the rate of photosynthesis and subsequently the rate at which this maximum storage capacity is reached, then leaf senescence will be advanced (Fu *et al*. 2014; Zani *et al*. 2020). This is also evidenced by the fact that trees that store nonstructural carbohydrates faster show earlier leaf senescence (Fu *et al*. 2014).

Over the past decades, an increasing number of phenological networks have been established to understand the phenological responses to climate change. As the largest phenological database worldwide, Pan European Phenology (PEP725) network (www.pep725.eu) (Templ et al. 2018) holds more than 12 million ground phenological records across 256 plant species, most of which spanned the years from 1951 to 2015. However, PEP725 network is constrained to a relatively small spatial scale consisting mostly of sites located in Central Europe. In contrast, the extracted phenological metrics from PhenoCam and remote-sensing products cover a large spatial scale, but only cover relatively short-term periods. In addition, eddy covariance technique has been widely applied to assess the photosynthetic carbon uptake and respiration of terrestrial ecosystems (Baldocchi et al. 2001). In particular, the FLUXNET (https://fluxnet.org/data/) provides a uniform and high-quality dataset of 212 eddy covariance flux towers worldwide. Therefore, it is important to combine different complementary datasets to provide a comprehensive understanding of the climatic response of autumn leaf senescence under global warming.

Using 500,000 phenological records for 15 temperate trees at 5,000 sites, phenological metrics derived from PhenoCam and satellite imagery, and 72 sites from FLUXNET network in the Northern Hemisphere (Fig. 1), we carried out detailed analyses of the responses of leaf senescence to warming and aim to disentangle the mechanisms underlying the observed contrasts in the responses of leaf senescence to warming and provide an ecological basis for predicting the trajectory of leaf senescence under future warming. We raise the hypothesis that the timing of leaf senescence is driven by both carbon sink limitation and nutrient resorption. Thus, the timing of leaf senescence is advanced by warmer temperatures in spring and summer, which lead to the carbon storage capacity being filled more rapidly, but is delayed by warmer temperatures in autumn, as a result of an extension in the remobilization of nutrients from old leaves for overwintering. Responses to past climate warming can provide a direct cue as to the likely trajectory of leaf senescence in the future.

**Fig. 1.**
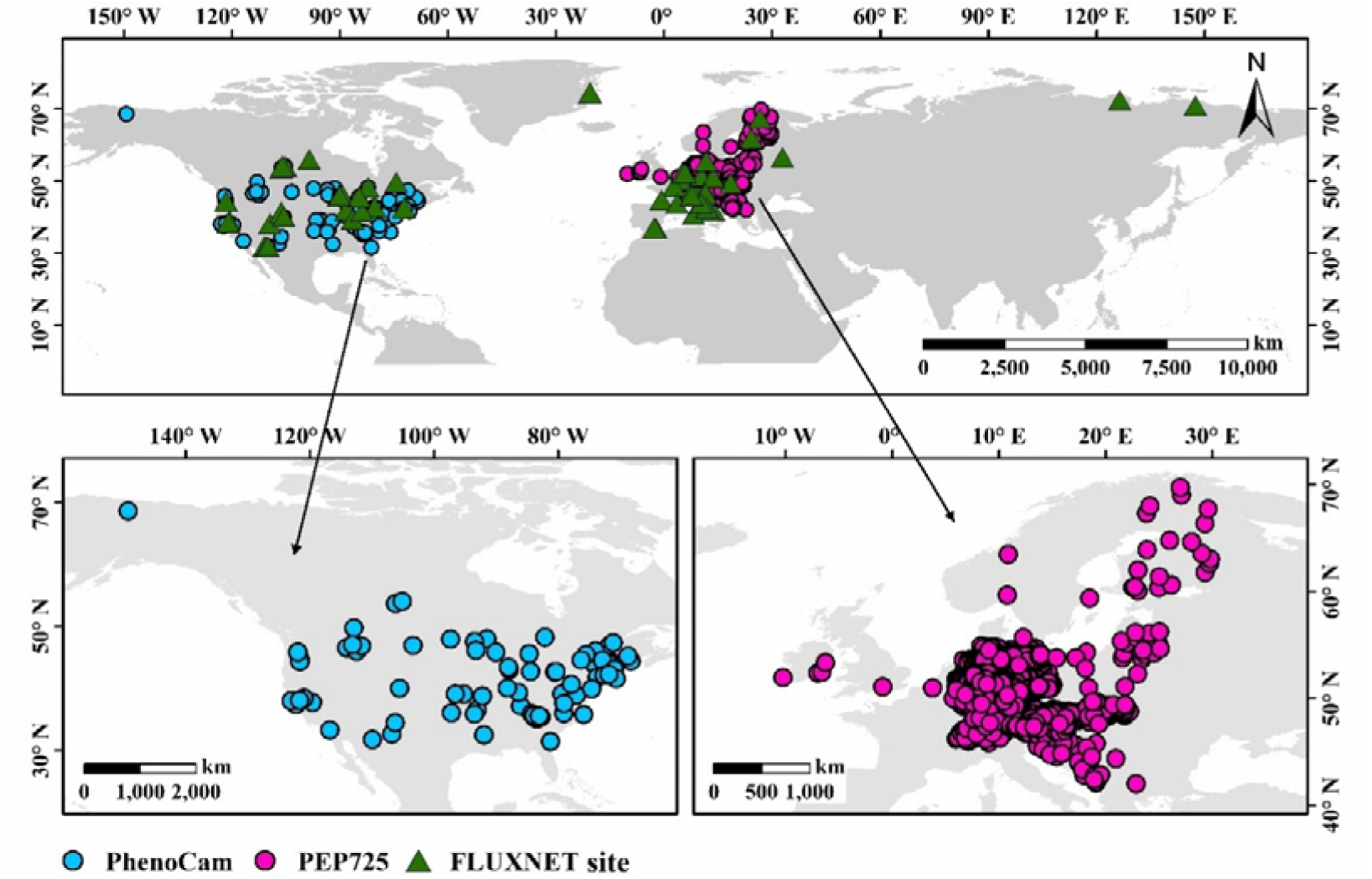
Locations of the phenological and FLUXNET sites used in this study. Phenological sites includes 5000 sites across central Europe selected from the Pan European Phenology (PEP725) database and 97 sites located in North America obtained from the PhenoCam network. A total 72 flux sites (>30°N) from the FLUXNET2015 dataset was selected.

## MATERIAL AND METHODS

### PEP725 phenological network

Ground observation phenological data were obtained from the Pan European Phenology (PEP725) network (www.pep725.eu) (Templ *et al*. 2018), one of the largest phenological database worldwide, which provides open-access *in situ* phenology observations in Europe collected by citizen scientists and researchers for science, research and education. The PEP725 network holds more than 12 million ground phenological records of 46 growth stages across 256 plant species and cultivars at nearly 20, 000 sites across 30 countries in Europe, with a majority of the sites being located in Germany. The phenological stages were defined according to the BBCH (Biologische Bundesanstalt, Bundessortenamt und CHemische Industrie) code (Meier 2001). Although the first phenological record dated back to 1868, most phenological observations were collected after 1951, the year when the plant phenology network was lunched in Europe. In the PEP725 network, leaf senescence was coded as 94 (BBCH). The date of leaf senescence is expressed as the day of year (DOY), which was defined as autumn coloring of leaves (50%). To identify and exclude outliers, median absolute deviation (MAD) method was used to filter the records of leaf senescence (Leys *et al*. 2013). Using a conservative criterion, we removed phenological records deviating by more than 2.5 times MAD (Leys *et al*. 2013). Then we selected 500,000 records of leaf senescence for 15 temperate species (Table S1) at 5,000 sites with at least 10 years of data between 1951 and 2015 across central Europe (Fig. 1). In addition, the corresponding records of leaf unfolding for these 15 species between 1951 and 2015 at each site were collected to determine the start of the growing season.

### PhenoCam network

Repeated photography from digital cameras set up at a fixed ground location has been widely applied to characterize the temporal changes in vegetation phenology in recent decades (Brown *et al*. 2016; Richardson *et al*. 2018a). The PhenoCam network is the largest cooperative database of digital phenocamera imagery. The network provides the dates of phenological transitions between 2000 and 2018 across different biomes in North America (Seyednasrollah *et al*. 2019). In the PhenoCam network, the 50^th^, 75^th^ and 90^th^ percentiles of the Green Chromatic Coordinate (G_CC_) were calculated to extract the dates of increase and decrease in greenness. The formula for G_CC_ is as follows:

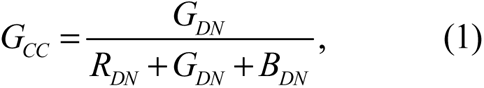

where *R_DN_*, *G_DN_* and *B_DN_* are, respectively, the average red, green and blue digital numbers (DN) across the region of interest. Previous studies have shown that the 90^th^ percentile of the GCC is effective at minimizing day-to-day variation due to weather conditions (e.g. clouds and aerosols) and illumination patterns (Sonnentag *et al*. 2012). Thus, we used the date on which the 90^th^ percentile GCC was observed to represent the date of leaf senescence.

### MODIS phenology product

The MODIS land surface phenology product (Collection 6 MCD12Q2) provides annual characteristics of vegetation phenology at a spatial resolution of 500 m between 2001 and 2017 on a global scale (Friedl *et al*. 2019). The phenological metrics were derived from the 8-day Enhanced Vegetation Index (EVI), which is calculated using MODIS nadir BRDF adjusted surface reflectances (NBAR-EVI2). Using this product, penalized cubic smoothing splines were used to fit the 8-day EVI time series and extract the onset of senescence, which was defined as the date when the fitted NBAR-EVI2 time series last crossed the 90^th^ percentile of the seasonal amplitude. The MCD12Q2 product was downloaded from the Land Processes Distributed Active Archive Center (LPDAAC) (https://lpdaac.usgs.gov/). In contrast to temperate and boreal regions, seasonal variations in vegetation dynamics are unclear in tropical and subtropical regions. We therefore excluded tropical and subtropical regions based on a map of terrestrial ecoregions worldwide (Dinerstein *et al*. 2017). Furthermore, we excluded cropland, as well as permanent snow and ice regions, based on the MODIS Landover classification product (MCD12Q1 version 6). The remaining biomes included Tundra, Boreal Forests/Taiga, Temperate Conifer Forests, Temperate Grasslands, Savannas & Shrublands, Mediterranean Forests, Woodlands & Scrub, Deserts & Xeric Shrublands, Temperate Broadleaf & Mixed Forests, Montane Grasslands & Shrublands.

### Climate data

Gridded daily mean (T_mean_), maximum (T_max_) and minimum (T_min_) temperatures, precipitation, radiation and humidity data with a spatial resolution of 0.25° in Europe were collected from the database E-OBS (http://ensembles-eu.metoffice.com). The period for temperature and precipitation data spans between 1951 and 2015, while radiation and humidity data were available between 1980 and 2015. Gridded CLM/ERAi soil moisture (0-10cm) data between 1980 and 2015 were downloaded from KNMI Climate Explorer (http://climexp.knmi.nl/select.cgi?id=someone@somewhere&field=clm_era_soil01). Global monthly mean temperature data with 0.5° spatial resolution between 2001 and 2018 were downloaded from the Climate Research Unit (CRU TS v4.04, https://crudata.uea.ac.uk/cru/data/hrg/cru_ts_4.04/cruts.2004151855.v4.04/). The E-OBS and CLM/ERAi climate datasets was used to analyze the effect of temperature on leaf senescence recorded *in situ* obtained from the PEP725 database. The CRU climate dataset was applied to analyze the effect of climate on the leaf senescence metrics extracted from the PhenoCam network and the MODIS vegetation phenology (MCD12Q2) product. The bilinear interpolation method was used to extract the climate data of all sites using the “raster” package (Hijmans *et al*. 2015) in R version 3.6.1 (R Core Team 2018).

The phenological records from the PEP725 database, which spanned the years from 1951 to 2015, covered a much longer period than those from the PhenoCam and MODIS datasets (only available since 2000). In addition, the PEP725 data were relatively more reliable than phenocam- and satellite-derived phenology because its leaf senescence records are taken manually *in situ*. The long-term gridded daily climate data in Europe obtained from the E-OBS database can be used to calculate climate index (e.g., growing degree-days) and further clarify the mechanisms underlying the climatic responses of leaf senescence. We were therefore able to test our hypotheses most directly using the PEP725 network and the corresponding E-OBS climate dataset. The PhenoCam and MODIS phenology products were used to test the robustness and generality of the results obtained from the PEP725 network in our study.

### Flux data

The flux dataset was download from FLUXNET (https://fluxnet.org/data/). The FLUXNET is a uniform and high-quality dataset based on regional flux networks worldwide. The FLUXNET2015 dataset (the latest released version) was downloaded from http://fluxnet.fluxdata. org/data/fluxnet2015-dataset/, which provides data on the exchange of carbon, water and energy of 212 sites across the globe, including over 1500 site-years, most of time series spanned between 2000 and 2014 (Pastorello *et al*. 2020). The FLUXNET2015 dataset has been processed using a uniform pipeline to reduce the uncertainty and improve the consistency across different sites (Pastorello *et al*. 2020), which has been widely applied to study the impact of climate change on carbon cycling in terrestrial ecosystems (Liu *et al*. 2019; Banbury Morgan *et al*. 2021). Due to the unclear vegetation carbon dynamics in tropical and subtropical regions, we only selected 72 sites (>□30°□N) across four vegetation types: Forest, Shrub, Grassland and Savanna in the Northern Hemisphere (Fig. 1).

### Temperature sensitivities

Temperature sensitivity (S_T_, change in days per degree Celsius) is expressed as the slope of a linear regression between the dates of phenological events and the temperature. This approach has been widely applied to assess phenological responses to global climate warming (Fu *et al*. 2015; Güsewell *et al*. 2017; Keenan *et al*. 2020). The S_T_ of leaf senescence was therefore used to investigate the effects of temperature during the growing season on leaf senescence. The length of growing season was defined as the period between the dates of leaf unfolding and leaf senescence for each species at each site. Using the daily climate data, we calculated the weekly and monthly mean temperature during the entire season for each species at each site. Then linear regression models were used to calculate the daily, weekly, and monthly S_T_ of leaf senescence throughout the entire season for each species at each site. The linear regression model was as follows:

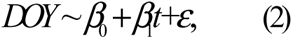

where *DOY* represents the date of leaf senescence; *t* represents the daily, weekly or monthly mean temperature; *β*_0_ is the intercept, *β*_1_ represent the S_T_ of leaf senescence; ε is the error of the model. In order to compare the effect of temperature on leaf senescence for different species at different sites, normalized anomalies (relative to the average) of temperature and leaf senescence dates were used in the linear regressions to calculate the S_T_ of leaf senescence for each species at each site (Chen *et al*. 2020; Keenan *et al*. 2020).

The mean dates of leaf unfolding and leaf senescence of the 15 studied species across the selected 5,000 sites from the PEP725 network were DOY 120 and DOY 280. In this context, we mainly considered daily, weekly and monthly S_T_ of leaf senescence from May to September. We applied linear regressions to test the temporal changes in the daily and weekly S_T_ of leaf senescence. One-way analysis of variance (ANOVA) followed by a Tukey’s HSD (honestly significant difference) test was used to examine differences in the monthly S_T_ of leaf senescence among months. From the calculated daily, weekly and monthly S_T_ of leaf senescence, we found that responses of leaf senescence to warming changed from negative in May and June to positive between July and September. We therefore divided the entire season into two periods: early season (May-June) and late season (July-September), and further calculated the mean S_T_ during the two periods to obtain the S_T_ during the early (S_T-Early_) and late (S_T-Late_) season, respectively. The sum of S_T-Early_ and S_T-Late_ of leaf senescence was used to represent the overall warming responses of leaf senescence during the whole season while leaves were present.

In addition, linear mixed models (Zuur *et al*. 2009) were used to pool all the data across species and sites and examined the overall S_T_ of leaf senescence during the early and late season. In the models, the response variable was leaf senescence date, the fixed effect was mean temperature during the early or late season, with species and site included as random intercept terms.

We followed Fu *et al*. (2015) to assess the effects of past climate warming on tree phenology. First, we calculated the mean temperature during the entire season (May-September) across all the 5,000 sites in Europe from 1951 to 2015. Using a 20-year smoothing window, we then identified the coldest and warmest periods: 1953-1972 and 1992-2011. The mean temperatures across the entire season during 1953-1972 and 1992-2011 were 14.57±0.61 and 15.52±0.70 respectively. Finally, we calculated and compared the S_T_ of leaf senescence during the early, late and entire season between 1953-1972 and 1992-2011. One-way ANOVA was used to test the difference in the S_T_ of leaf senescence between the two periods.

To test the robustness and generality of results obtained from the PEP725 network, we further calculated the monthly S_T_ of leaf senescence between May and September during 2000-2018 based on the dates of leaf senescence extracted from the PhenoCam network and MODIS phenology product. Because seasonal cycles in cropland are considerably influenced by human activities, we first excluded those sites in cropland and selected 97 sites (Fig. 1) with at least 5 years of data from the PhenoCam network. Then we calculated the monthly S_T_ of leaf senescence between May and September during the period 2000-2018 for each site across North America. Because most of the selected sites (61 sites) were located in deciduous broadleaf forests, we did not address the difference in the S_T_ of leaf senescence among biomes using the PhenoCam network. Instead, we calculated and compared the monthly S_T_ of leaf senescence between May and September among biomes in the Northern Hemisphere based on the phenological metrics derived from the MODIS phenology product. In contrast to temperate and boreal regions, seasonal variations in vegetation dynamics are unclear in tropical and subtropical regions. We therefore excluded tropical and subtropical regions based on a map of terrestrial ecoregions worldwide (Dinerstein *et al*. 2017). Furthermore, we excluded cropland, as well as permanent snow and ice regions, based on the MODIS Landover classification product (MCD12Q1 version 6). The remaining biomes included Tundra, Boreal Forests/Taiga, Temperate Conifer Forests, Temperate Grasslands, Savannas & Shrublands, Mediterranean Forests, Woodlands & Scrub, Deserts & Xeric Shrublands, Temperate Broadleaf & Mixed Forests, Montane Grasslands & Shrublands. One-way ANOVA followed by a Tukey’s HSD test was used to test the difference in the monthly S_T_ of leaf senescence among biomes.

In addition to temperature, autumn phenology is also influenced by other environmental factors (Misson *et al*. 2011; Liu *et al*. 2016; Chen *et al*. 2020). Using partial correlation analysis, we excluded the covariate effects of soil moisture, precipitation, radiation, humidity and further examined the relationship between monthly mean temperature from May to September and leaf senescence dates. To test the effect of drought stress on leaf senescence, we also calculated the partial correlation coefficients between soil moisture and leaf senescence between May and September for each species at each site. Furthermore, we quantified and compared the relative influences of temperature, soil moisture, precipitation, radiation, humidity on leaf senescence date in the growing season using boosted regression trees (BRTs), an ensemble statistical learning method (Elith *et al*. 2008) that has been widely applied to ecological modeling and prediction (Chen *et al*. 2018; Davis *et al*. 2019; Lemm *et al*. 2021). We performed the BRTs using the GBM package (Ridgeway 2007) of R (R Core Team, 2018), where 10-fold cross validation was used to determine the optimal number of iterations. Because gridded soil moisture, solar radiation, humidity data was only available since 1980, the observations between 1980 and 2015 from the PEP725 network were selected for the multiple factor analysis.

### Effect of growing degree-days on leaf senescence

Using the daily temperature from the E-OBS database, we calculated the accumulated growing degree-days (GDDs) in each month during the growing season from May to September at each site used in the PEP725 dataset. The base temperatures were set as 5 when calculating the GDDs. Because the temperature responses of leaf senescence changed from negative in May and June to positive between July and September, we divided the entire growing season into two periods, early growing season (May-June) and late growing season (July-September), and calculated the mean accumulated GDDs during each of the two periods. Then linear regression models were used to examine the effects of GDDs on the leaf senescence dates (change in days GDD^-1^) in years under low and high nighttime temperature conditions during the early and late growing season at each site selected from the PEP725 database. The classification of early (or late) seasons with low and high nighttime temperature was based on whether the mean daily T_min_ during the early (or late) growing season for a given year at a site was, respectively, below or above the long-term average during 1951-2015. One-way ANOVA was used to test for differences in the effect of GDDs on leaf senescence between low and high nighttime temperature conditions during the early and late growing seasons. Using the FLUXNET2015 data, we calculated and compared the differences in the nighttime respiration during the early season (May and June) and the number of frost days (T_min_ <0□) in late autumn (October and November) during years with low and high nighttime temperature using one-way ANOVA analysis. The classification of the seasons in years with low and high nighttime temperature was based on whether the mean daily T_min_ during the early (or late) growing season for a given year at a site was, respectively, below or above the long-term average during 2000-2014.

### Photosynthetic carbon assimilation

Using the FLUXNET2015 dataset, we examined the temporal changes in the photosynthesis carbon assimilation during the growing season based on the Net Ecosystem Exchange (NEE). The NEE measures the net carbon exchange between ecosystem and and atmosphere, which approximately equals to net primary productivity (NPP) when soil respiration approaches zero, but with opposite sign (Chapin *et al*. 2006; Lasslop *et al*. 2010). In our study, the opposite NEE is therefore used to estimate net photosynthetic carbon assimilation. Singular Spectrum Analysis (SSA) was applied to smooth the daily NEE of each year at each site between 2000 and 2014 to minimize the noise. One-way ANOVA analysis was used to compare the net carbon assimilation during the early season (before June) and late season (after June).

All data analyses were conducted using R version 3.6.1 (R Core Team 2018).

## RESULTS

Using records of leaf senescence for 15 temperature tree species at 5,000 sites from the PEP725 network, we found the mean S_T_ of leaf senescence was negative in May and June, while it gradually converted to positive between July and September (Fig. 2a). This suggested that increasing temperatures during early season advanced leaf senescence, but increasing temperatures during the late season delayed leaf senescence (see for example, *Fagus sylvatica* and *Quercus robur* in Figs S1 and S2). In addition, the delaying effects of temperature on leaf senescence started from July generally showed an increasing trend, reaching a maximum in September (Fig. 2a). Based on the daily and weekly S_T_ of leaf senescence, we also observed a significant increase in S_T_ throughout the entire season (Fig. S3).

**Fig. 2.**
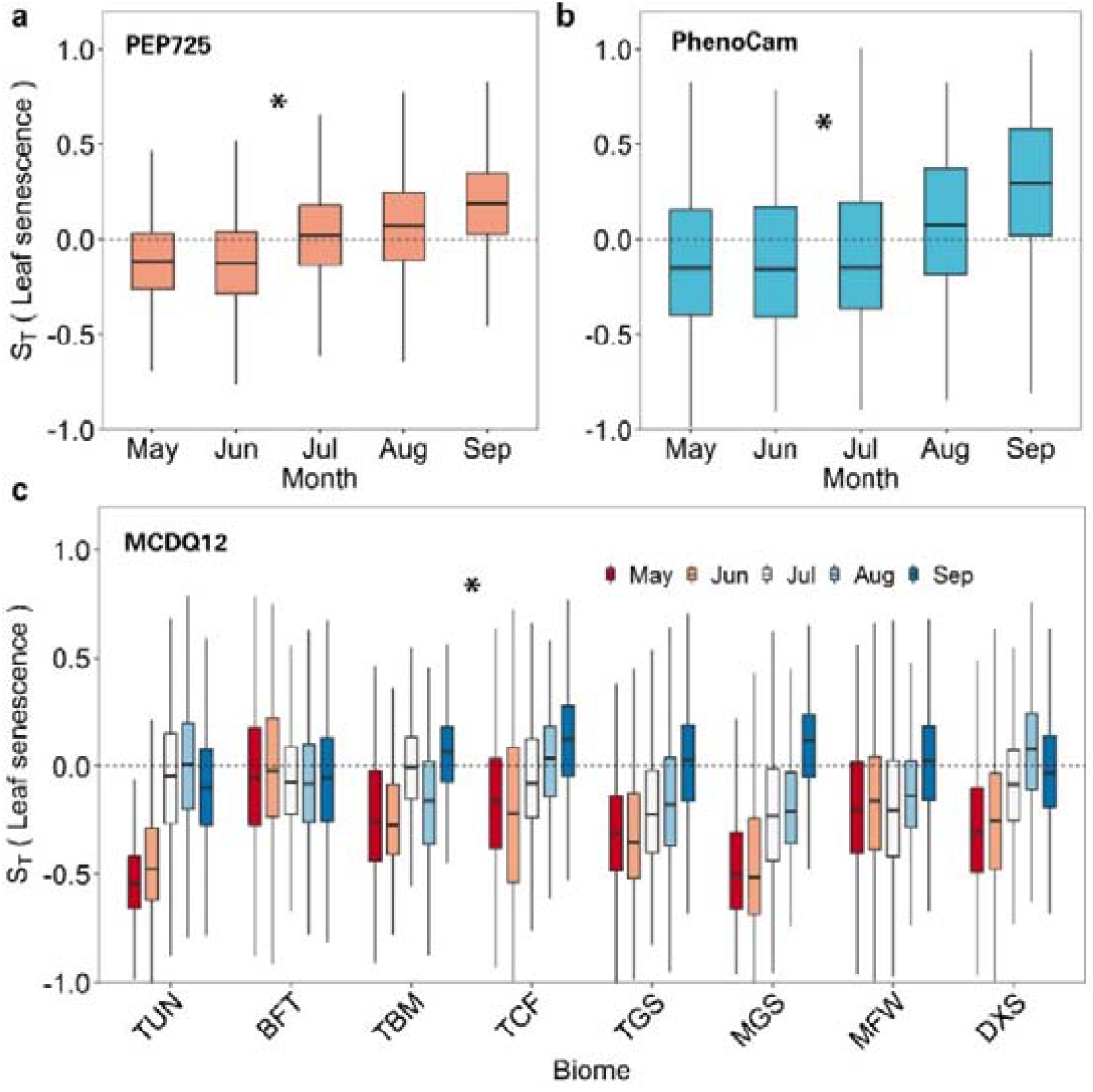
Temperature sensitivities (S_T_, change in days ^−1^) of leaf senescence during the growing season between May and September. The calculated S_T_ was based on **(a)** records of leaf senescence for 15 temperate tree species at 5,000 sites in Europe, and phenological metrics extracted from **(b)** the PhenoCam network and **(c)** the MODIS phenology product (MCD12Q2 version 6) for different biomes. The length of each box indicates the interquartile range, the horizontal line inside each box the median, and the bottom and top of the box the first and third quartiles respectively. The biomes are Tundra (TUN), Boreal Forests/Taiga (BFT), Temperate Broadleaf & Mixed Forests (TBM), Temperate Conifer Forests (TCF), Temperate Grasslands, Savannas & Shrublands (TGS), Montane Grasslands & Shrublands (MGS), Mediterranean Forests, Woodlands & Scrub (MFW), and Deserts & Xeric Shrublands (DXS). The asterisks indicate a significant difference in the S_T_ during the early (May-June) and the late (July-September) season (*P*<0.05).

According to the linear mixed models, the overall S_T_ of leaf senescence during the early and late season across all species and sites were approximately −1.14 and +1.33 days per degree Celsius, while S_T_ of leaf senescence of the total season was +0.12 days per degree Celsius (Table S2). During the early season, the monthly S_T_ of leaf senescence between May and June was similar (Table S2). During the late season, the monthly S_T_ of leaf senescence in September was the strongest among all the months (Table S2). The absolute S_T_ of leaf senescence in September was stronger than that in May (Table S2).

Using the PhenoCam network, we again observed a transition in the S_T_ of leaf senescence in North America from May to September (Fig. 2b). The effect of temperature on leaf senescence was negative in May and July (Fig. 2b). However, a positive effect was observed in August and September in North America (Fig. 2b), confirming the results from the PEP725 network.

Based on phenology metrics extracted from MODIS, we consistently observed a transition in the S_T_ of leaf senescence from May to September across all biomes except deserts and xeric shrublands in the Northern Hemisphere (Fig. 2c). In May and June, the effects of temperature on leaf senescence were negative across temperate and boreal biomes (Fig. 2c). We also observed negative effects of temperatures in May and June on leaf senescence in tundra, alpine and Mediterranean regions (Fig. 2c). The effects of temperature gradually became positive in August or September in these biomes (Fig. 2c). For deserts and xeric shrublands, we similarly observed a negative effect of temperature on leaf senescence in May and June (Fig. 2c). These negative effects were significantly weaker in deserts and xeric shrublands compared to other biomes (Fig. 2c). However, the effects of temperature remained negative throughout the growing season in deserts and xeric shrublands (Fig. 2c). In these environments, the negative effect of temperature on leaf senescence was stronger in August than in June (Fig. 2c). When we mapped the monthly S_T_ of leaf senescence during the growing season, we also observed a transition in the S_T_ of leaf senescence during the growing season in the Northern Hemisphere (Fig. S4). Overall, the S_T_ showed a significant increase from the early to the late season, as indicated by the larger S_T_ in September than in May (Fig. S4f).

After excluding the effects of other climate variables, using partial correlation analysis we also observed a negative response of leaf senescence to temperature during the early season, but a positive response during the late season (Fig. S5). In contrast to temperature, we observed no significant difference in the responses of leaf senescence to soil moisture (Fig. S6). A positive effect of soil moisture on leaf senescence in May and July was observed (Fig. S6). According to the calculated relative influence, temperature had the strongest effect on leaf senescence, followed by soil moisture and radiation (Fig. S7).

Using the FLUXNET2015 data, we detected an obvious changing point at the end of June (DOY 180) for the net daily carbon assimilation (Fig. 3). Generally, net carbon assimilation was positive during the early season (before June) but was negative during the late season (after June) (Fig. 3a). This suggested that carbon assimilation mainly occurs before June. The difference in the net carbon assimilation between early season (before June) and late season (after June) in forest was the largest, followed by grassland and savanna (Fig. 3b).

**Fig. 3.**
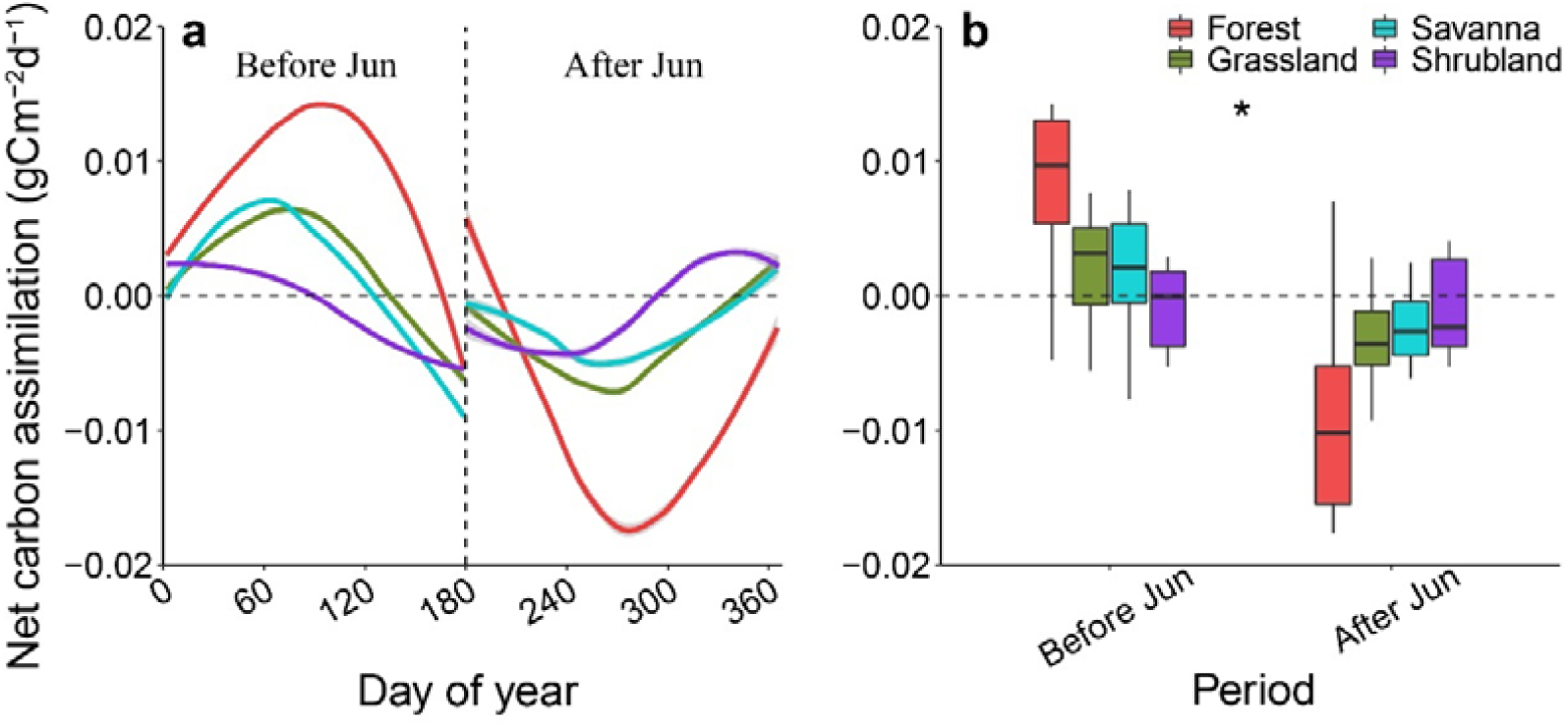
Change in the net daily photosynthetic carbon assimilation (g Cm^-2^d^-1^). The estimation of the net carbon assimilation was calculated by multiplying the Net Ecosystem Exchange (NEE) by −1. The length of each box in **(b)** indicates the interquartile range, the horizontal line inside each box the median, and the bottom and top of the box the first and third quartiles respectively. The asterisks indicate a significant difference in the net photosynthetic carbon assimilation during the early season (before June) and late season (after June). Different color lines and boxes represent different vegetation types. The dashed vertical line indicates the change point of net photosynthetic carbon assimilation (DOY 180).

Results showed that the timing of leaf senescence was also advanced by greater GDDs during the early season (May-June), but delayed by greater GDDs during the late season (July-September) (Fig. 4ab). We found that during the early season, the negative effect of GDDs on leaf senescence was stronger during years with low nighttime temperatures (*P*<0.001, Fig. 4a). In addition, during the late season the positive effect of GDDs on leaf senescence was weaker during years with low nighttime temperature (*P*<0.001, Fig. 4b). When nighttime temperature was higher, nighttime ecosystem respiration was significantly greater during the early season (Fig. 4c), while the number of frost days was significantly lower during the late season (Fig. 4d).

**Fig. 4.**
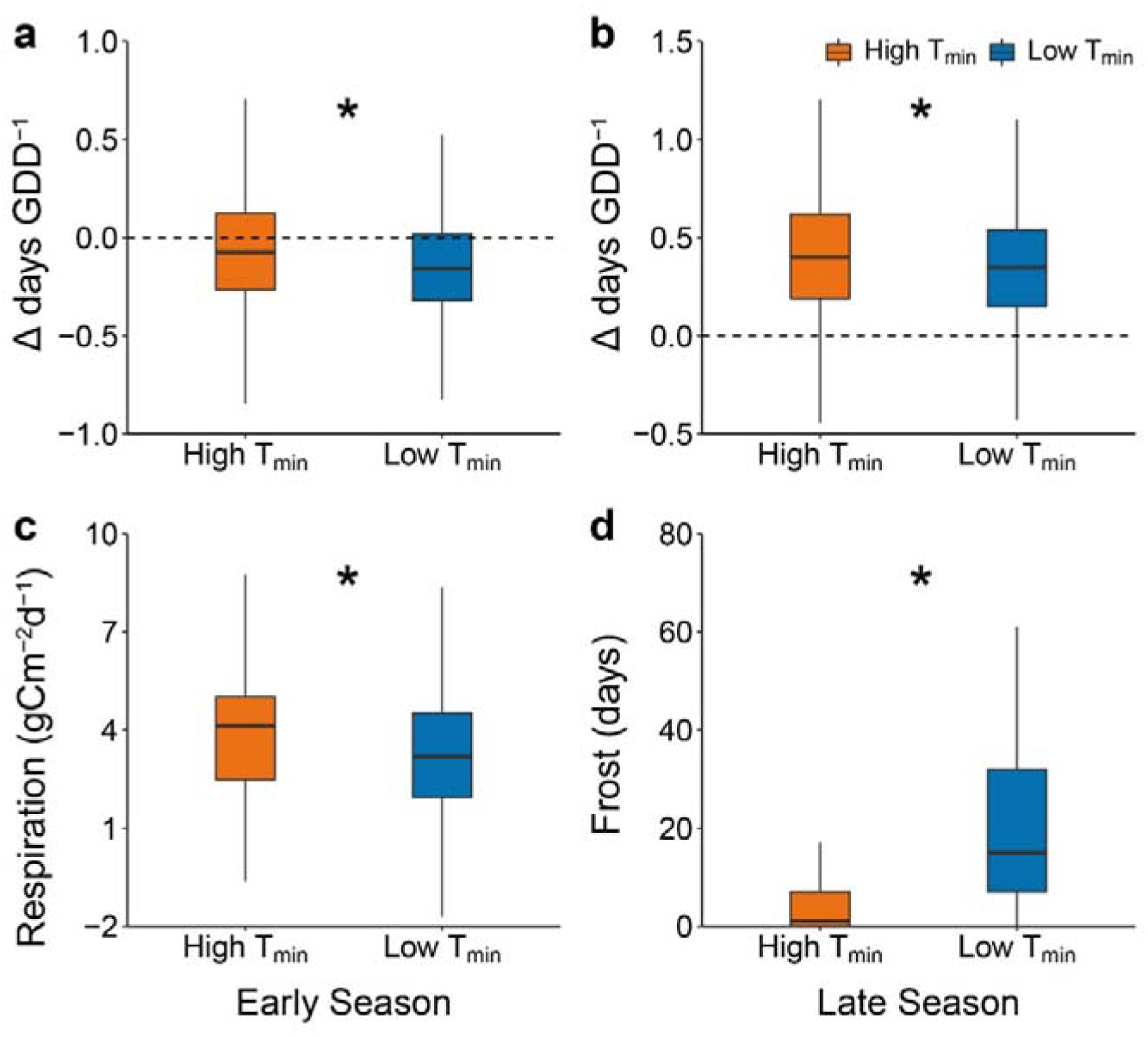
**(a, b)** Effects of growing degree-days (GDD) on leaf senescence during the early and late season in 15 temperate tree species at 5,000 sites in Europe between 1951 and 2015 using data from the Pan European Phenology (PEP725) network. (**c, d**) Difference in the nighttime respiration during the early season and frost days (Tmin < 0 □) in late autumn (October and November) using the FLUXNET data. The results are represented separately for seasons with low and high nighttime temperatures (daily minimum temperature, Tmin, □). The classification of the seasons was based on whether the mean daily Tmin during the (early or late) season for a given year was, respectively, below or above its long-term average. The length of each box indicates the interquartile range, the horizontal line inside each box the median, and the bottom and top of the box the first and third quartiles respectively. The asterisks indicate a significant difference in seasons with low and high nighttime temperature (*P*<0.05).

To assess the effects of climate warming on leaf senescence, we used the PEP725 dataset to calculate and compare the S_T_ of leaf senescence between the coldest and the warmest 20-year periods: 1953-1972 and 1992-2011. We found that both the S_T-Early_ and S_T-Late_ of leaf senescence were significantly higher during 1992-2011 than those during 1953-1972 (Fig. 5, *P*<0.05). However, S_T-Late_ of leaf senescence during 1992-2011 increased more compared to S_T-Early_ of leaf senescence (Fig. 5). This indicated that leaf senescence delayed more with the increasing temperatures during the late season during 1992-2011. For example, between 1953 and 1972 leaf senescence of *Fagus sylvatica* at several sites was advanced by increasing temperature during the late season, but was delayed by late season warming between 1992 and 2011 (Fig. S8). The S_T_ of leaf senescence during the whole growing season, i.e., the sum of S_T-Early_ and S_T-Late_ of leaf senescence, also showed a significant increase (Fig. 5, *P*<0.05).

**Fig. 5.**
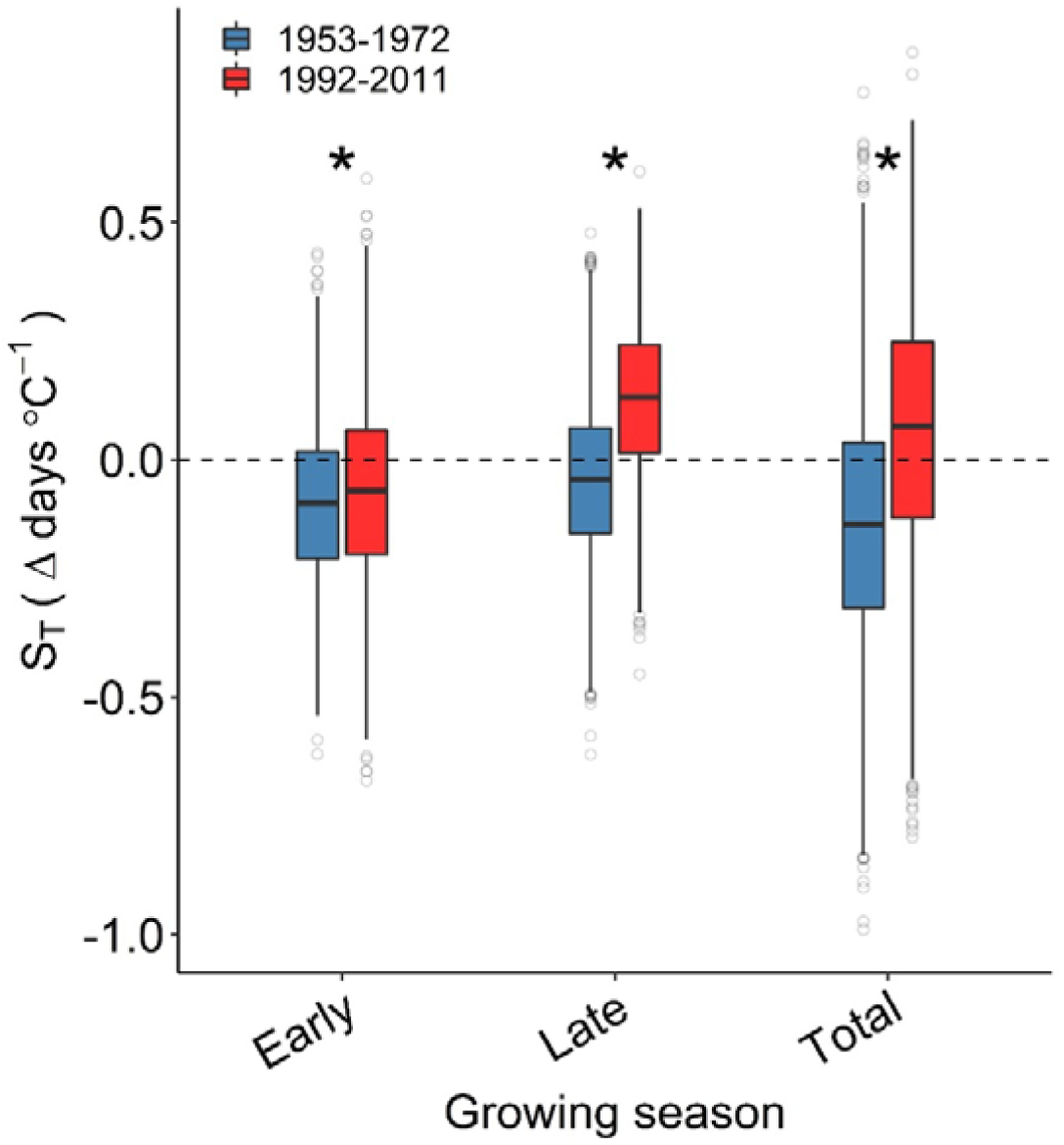
Temperature sensitivities (ST, change in days □ −1) of leaf senescence in 15 temperate tree species during 1953-1972 and 1992-2011. Calculations of the ST values were based on the temperature in early (May and June), late (July-September) and entire (May-September) growing seasons. The length of each box indicates the interquartile range, the horizontal line inside each box the median, and the bottom and top of the box the first and third quartiles respectively. The asterisks indicate a significant difference in the ST between 1953-1972 and 1992-2011 (*P*<0.05).

## DISCUSSION

Earlier leaf senescence reduces photosynthetic carbon assimilation and nutrient resorption efficiency (i.e. the proportion of nutrients resorbed from old leaves) (Estiarte & Penuelas 2015). However, trees experiencing late leaf senescence are more at risk from frost (Schwartz 2003; Hartmann *et al*. 2013), which may reduce nutrient resorption (Estiarte & Penuelas 2015). The optimal timing of leaf senescence is therefore likely to be a trade-off between photosynthetic carbon assimilation and autumnal nutrient resorption at different stages in the seasonal functioning of leaves (Keskitalo *et al*. 2005; Fracheboud *et al*. 2009) (Fig. 6). When trees reach their maximum carbon storage capacity, they will initialize nutrient resorption and senescence. Accordingly, more efficient accumulation of carbohydrates with warmer temperatures in the early season will result in a relatively shorter period being required to reach the maximum carbon storage capacity (Peng *et al*. 2013). Thus, early season warming advances leaf senescence. Using the flux data, we further found net carbon assimilation converted from positive to negative at the end of June (DOY 180). This provides direct physiological evidence that photosynthetic carbon assimilation mainly occurred during the early season (before June). However, warmer temperatures later in the season may reduce the risk of late autumn frost (Vitasse *et al*. 2014), enhancing the activities of photosynthetic enzymes (Shi *et al*. 2014) and reducing the rate of chlorophyll degradation (Fracheboud *et al*. 2009; Estiarte & Penuelas 2015). This may prolong nutrient remobilization from leaves, reduce degradation rate of organelle dismantling, increase leaf longevity and eventually delay the final stage of leaf senescence (Kikuzawa 1995). Therefore, leaf senescence was advanced by warming during the early season, but was delayed by the warming during the late season in temperate regions. However, leaf senescence was advanced by warming throughout both early and late seasons in deserts and xeric shrublands. This may result from drought stress caused by warmer autumns in arid regions increasing evaporative demand (Allen *et al*. 2010; Chen *et al*. 2017) and thus initiating leaf senescence early (Estiarte & Penuelas 2015; Wu *et al*. 2018; Chen *et al*. 2020), supported by the positive correlations between soil moisture and leaf senescence in May and July.

**Fig. 6.**
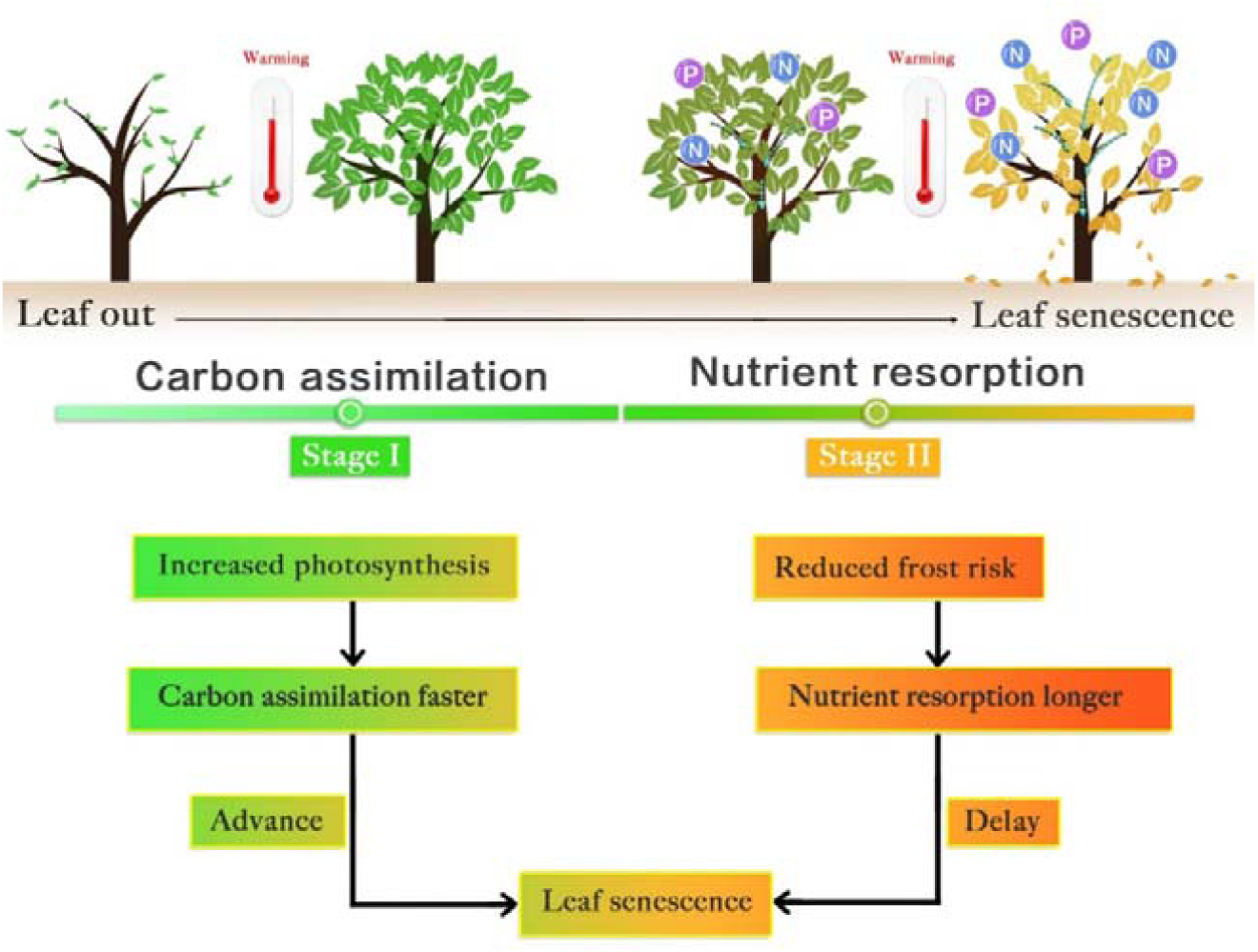
A schematic diagram of the contrasting responses of leaf senescence to warming during two seasonal stages of tree leaf development, in which photosynthetic carbon assimilation or nutrient resorption (of nitrogen, N and phosphorus, P) take place respectively.

The parameter growing degree-days (GDDs) only takes into account the heat accumulated above the minimum threshold of temperature that must be exceeded for tree growth (Briere *et al*. 1999; Miller *et al*. 2001) and therefore provides a more accurate physiological assessment of leaf senescence in response to warming (Wu *et al*. 2018; Chen *et al*. 2020). Our observation that leaf senescence was advanced by GDD-based warming during the early growing season, but was delayed by GDD-based warming during the late resorption season, are consistent with the contrasting warming responses of leaf senescence reported previously (Menzel *et al*. 2006; Jeong *et al*. 2011; Gill *et al*. 2015). These results similarly support trade-off of leaf senescence between carbon assimilation and nutrient resorption. We further examined this trade-off by comparing the warming responses of leaf senescence when the nighttime temperature changed, because of the asymmetric effects of nighttime temperature on carbon assimilation during the early growing season and frost avoidance during the late resorption season (Peng *et al*. 2013; Chen *et al*. 2020). In particular, accumulation of carbohydrates is likely to be more efficient when nighttime temperature is low, due to reduced nighttime respiration (Peng *et al*. 2013). As a result, trees will reach their maximum carbon capacity quickly when nighttime temperatures are lower according to such a trade-off assumption. By contrast, trees can be expected to accomplish nutrient resorption rapidly during the late season in order to reduce the risk of frost when nighttime temperatures are lower (Silvestro *et al*. 2019). Consistent with this, we found that the signals of the effects of warming on leaf senescence were stronger or weaker when the nighttime temperature was lower during the early or late season respectively. All of these findings indicate that not only the carbon sink limitation in the early season but also nutrient resorption in the late season should be considered when modelling autumn phenology of temperate trees under future warming scenarios. The results of warming modelling based mainly on sink limitation in the early season predict advancing of leaf senescence (Zani *et al*. 2020). However, our statistical analyses of leaf senescence during the warmest 20-year periods across both seasons suggest that global warming may delay leaf senescence in the future. Nonetheless, seasonal differences in the responses to warming need to be considered when modelling autumn phenology and carbon cycling.

In addition to temperature, photoperiod may influence tree phenology (Körner & Basler 2010). As photoperiod remains unchanged across years for a given location, a relatively conservative climatic response is therefore expected when trees rely on the photoperiod to determine phenology (Basler & Körner 2012; Way & Montgomery 2015; Flynn & Wolkovich 2018). Compared with spring leaf out, leaf senescence has been reported to show a more conservative warming response (Menzel *et al*. 2006). For this reason, autumnal phenological events are commonly considered to be more sensitive to photoperiod compared with spring events (Way & Montgomery 2015). However, such a conservative response to warming may be due to the counterbalancing effects of warming on leaf senescence at different seasons. Additionally, despite the photoperiodic control of leaf senescence (Way & Montgomery 2015; Singh *et al*. 2017), we found that temperature alone had strong predictive power even when photoperiod was not considered, indicating the dominant role of temperature in leaf senescence.

Overall, our findings based on three large and complementary datasets illustrate that the onset of leaf senescence is advanced under early season warming, but delayed when warming occurs in the late stages of the growing season. Although further controlled warming experiments in different seasons should be conducted to test the contrasting seasonal climatic responses of leaf senescence, our study provides new insights into how to accurately predict whether leaf senescence will be delayed or advanced in response to climate warming (Menzel *et al*. 2006). If future warming spans both early and late seasons in temperate regions, as found previously (Menzel *et al*. 2006; Gill *et al*. 2015), leaf senescence could be delayed, rather than advanced.

## ACKNOWLEDGEMENTS

We acknowledge all members of the PEP725 network for collecting and providing the phenological data. This research was supported by the Strategic Priority Research Program of the Chinese Academy of Sciences (XDB31010300) and the National Key Research and Development Program of China (2017YFC0505203), and also by the National Natural Science Foundation of China (grant numbers 31590821, 91731301 and 31561123001), the Fundamental Research Funds for the Central Universities (2018CDDY-S02-SCU and SCU2019D013), and National High-Level Talents Special Support Plans.

## AUTHOR CONTRIBUTIONS

LC and JL designed the research. LC performed the data analysis. LC wrote the paper with the inputs of SR, NGS and JL. All authors contributed to the interpretation of the results and approved the final manuscript.

## DATA ACCESSIBILITY

The ground observation phenological data are available at www.pep725.eu. The phenological metrics derived from digital camera imagery are available at https://lpdaac.usgs.gov/. The phenology data extracted from satellite images can be downloaded from https://daac.ornl.gov/cgi-bin/dsviewer.pl?ds_id=1674. The climate data used in this study are available at http://ensembles-eu.metoffice.com and https://crudata.uea.ac.uk /cru/data/hrg/ cru_ts_4.04/cruts.2004151855.v4.04/.

## SUPPORTING INFORMATION

**Table S1.** Dates of leaf senescence of the15 temperate species selected from the PEP725 phenological network. For each species, mean dates of leaf senescence (Mean), lower and upper limit of the 95% confidence interval (CI), and standard deviation (SD) of leaf senescence dates were listed.

| Species | Latin name | Common name | Mean | SD | Lower<br>(95%CI) | Upper<br>(95%CI) |
| --- | --- | --- | --- | --- | --- | --- |
| 1 | *Aesculus hippocastanum* L. | Horse chestnut | 277.0 | 11.7 | 276.9 | 277.0 |
| 2 | *Betula pendula* Roth | Silver birch | 278.9 | 12.8 | 278.9 | 279.0 |
| 3 | *Fagus sylvatica* L. | European beech | 282.6 | 12.1 | 282.5 | 282.7 |
| 4 | *Quercus robur* L. | English oak | 287.9 | 12.1 | 287.9 | 288.0 |
| 5 | *Prunus avium* (L.) L. | Sweet cherry | 284.7 | 13.9 | 284.5 | 284.9 |
| 6 | *Tilia cordata* Mill. | Lime | 283.6 | 13.8 | 283.1 | 284.0 |
| 7 | *Acer platanoides* L. | Norway maple | 263.0 | 9.5 | 262.0 | 263.9 |
| 8 | *Prunus domestica* L. | Common plum | 288.7 | 12.2 | 287.2 | 290.2 |
| 9 | *Larix decidua* Mill. | European larch | 294.1 | 12.8 | 293.9 | 294.2 |
| 10 | *Vitis vinifera* L. | Grape vine | 284.9 | 10.7 | 284.5 | 285.4 |
| 11 | *Malus domestica* Borkh. | Apple | 279.7 | 12.7 | 278.5 | 281.0 |
| 12 | *Corylus avellana* L. | Common hazel | 280.6 | 13.4 | 279.8 | 281.4 |
| 13 | *Sorbus aucuparia* L. | Rowan | 274.3 | 12.6 | 273.9 | 274.8 |
| 14 | *Betula pubescens* Ehrh. | White birch | 282.8 | 15.1 | 281.9 | 283.7 |
| 15 | *Populus tremula* L. | European aspen | 271.0 | 14.5 | 269.5 | 272.4 |

**Table S2.** Results of the linear mixed models for the overall temperature sensitivity (S_T_, change in days per degree Celsius) of leaf senescence during the early (May-June) and late season (July-September) across all species and site.

| Season | Period | $S_T$ | SE | $t$ value | $P$ value |
| --- | --- | --- | --- | --- | --- |
| Early | May | -0.87 | 0.01 | -63.31 | < 0.001 |
|  | Jun | -0.86 | 0.02 | -53.74 | < 0.001 |
|  | May-Jun | -1.24 | 0.02 | -70.34 | < 0.001 |
| Late | Jul | 0.42 | 0.01 | 33.90 | < 0.001 |
|  | Aug | 0.21 | 0.02 | 13.45 | < 0.001 |
|  | Sep | 1.08 | 0.01 | 75.1 | < 0.001 |
|  | Jul-Sep | 1.33 | 0.02 | 62.63 | < 0.001 |
| Total | May-Sep | 0.12 | 0.03 | 4.69 | < 0.001 |

**Fig. S1.**
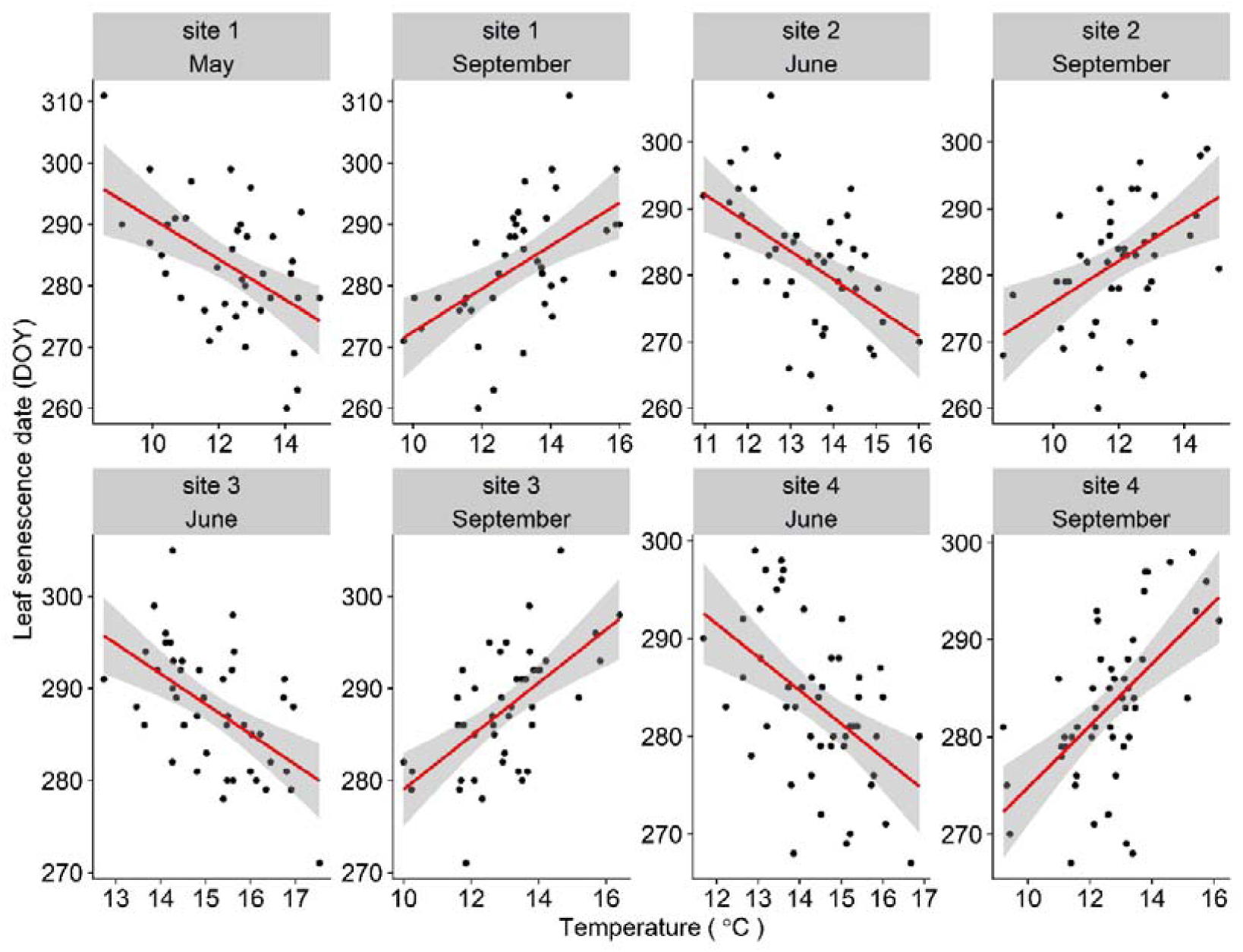
Effects of monthly mean temperature during the early (May and June) and late (July-September) on leaf senescence dates of *Fagus sylvatica* at several sites selected from the Pan European Phenology (PEP725) database. The leaf senescence date was expressed as the day of year (DOY). The shaded area indicates the 95% confidence intervals of the fitted regression lines.

**Fig. S2.**
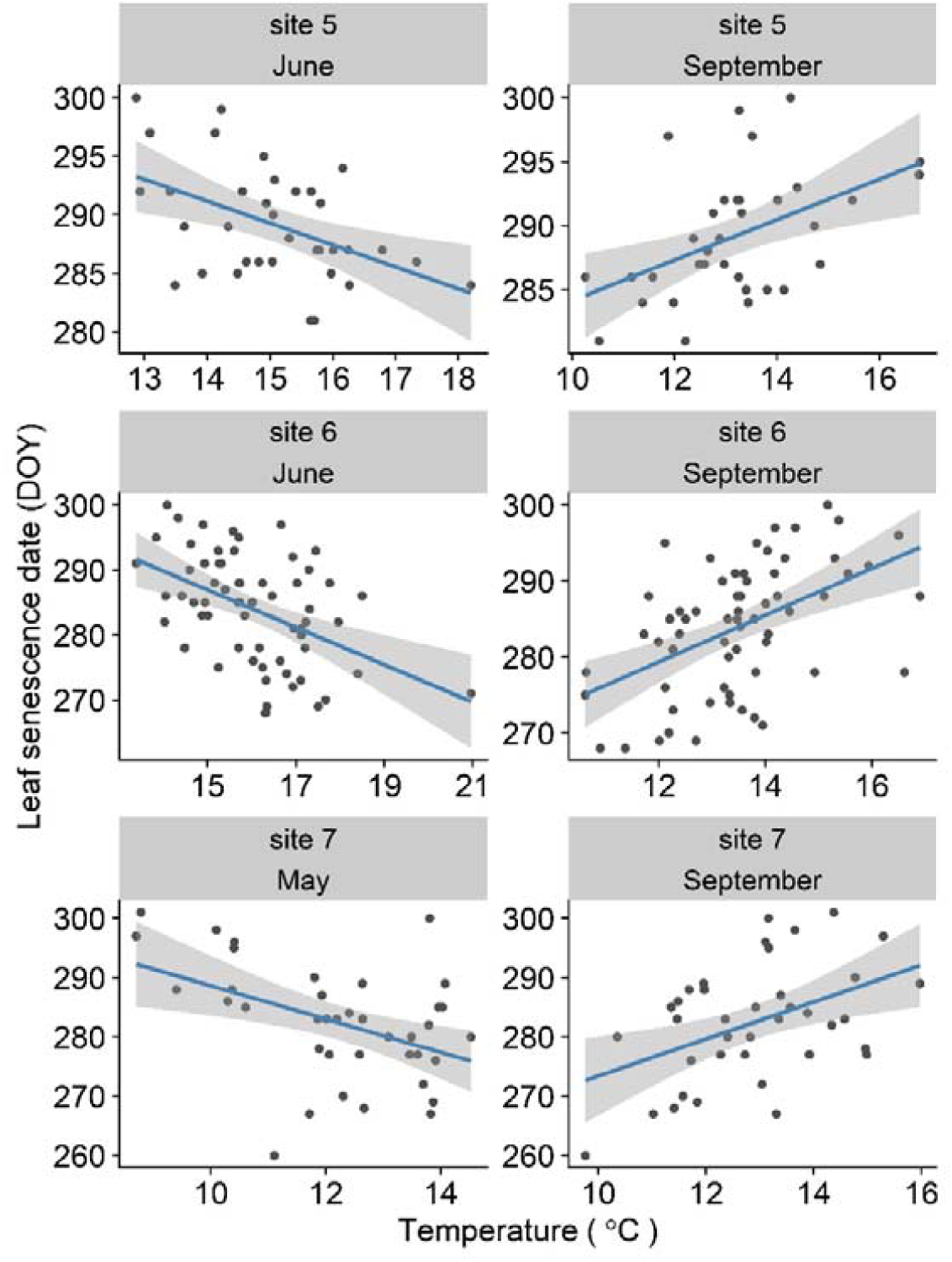
Effects of monthly mean temperature during the early (May and June) and late (July-September) on leaf senescence dates of *Quercus robur* at several sites selected from the Pan European Phenology (PEP725) database. The leaf senescence date was expressed as the day of year (DOY). The shaded area indicates the 95% confidence intervals of the fitted regression lines.

**Fig. S3.**
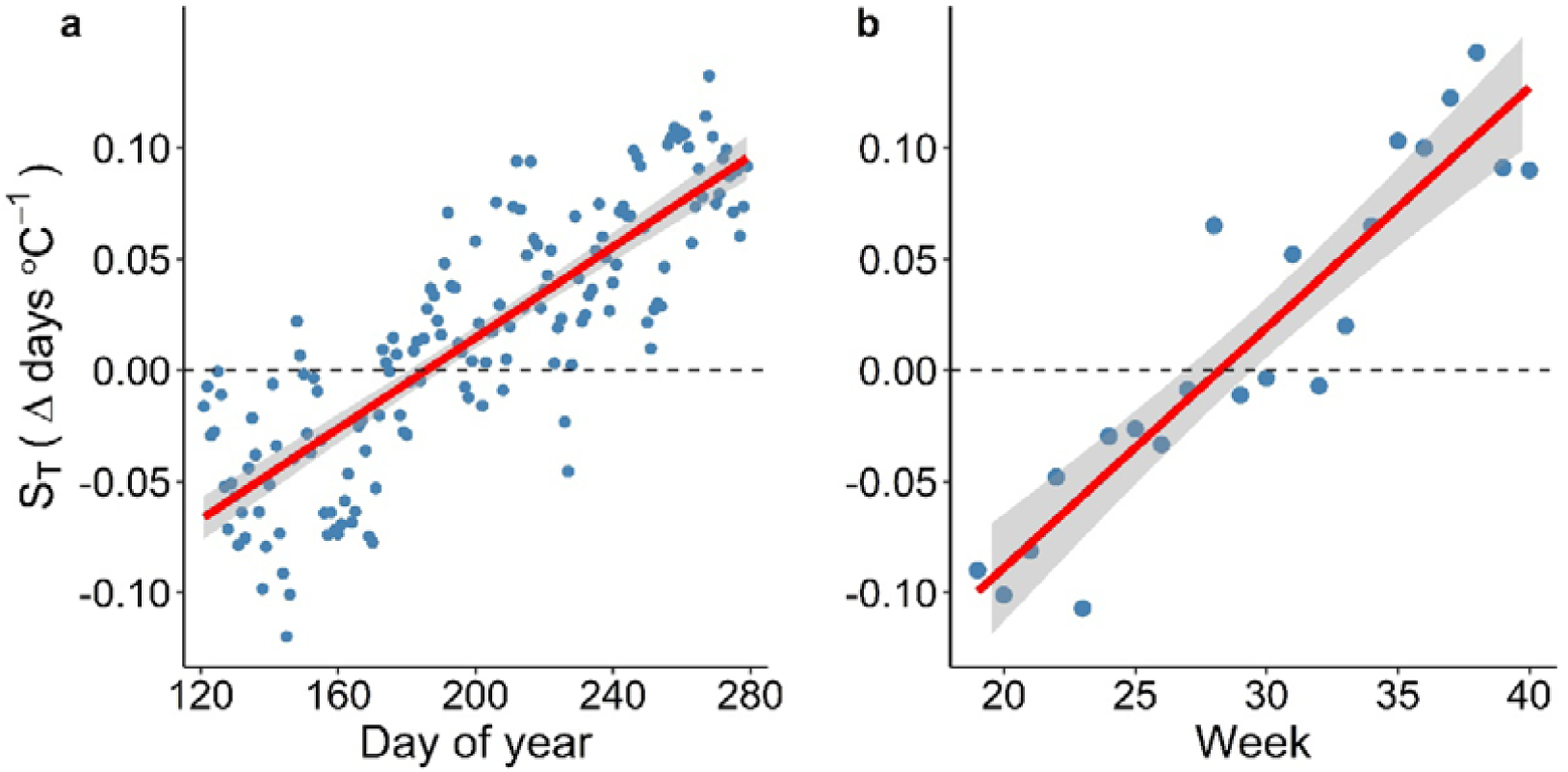
Daily and weekly temperature sensitivities (ST, change in days □ −1) of leaf senescence during the growing season between May and September. The calculated S_T_ values were based on records of leaf senescence for 15 temperate tree species at 5,000 sites in Europe. Each point represents the mean daily or weekly S_T_ of leaf senescence calculated across all 15 species at 5000 sites. The shaded areas indicate the 95% confidence intervals of the fitted regression lines.

**Fig. S4.**
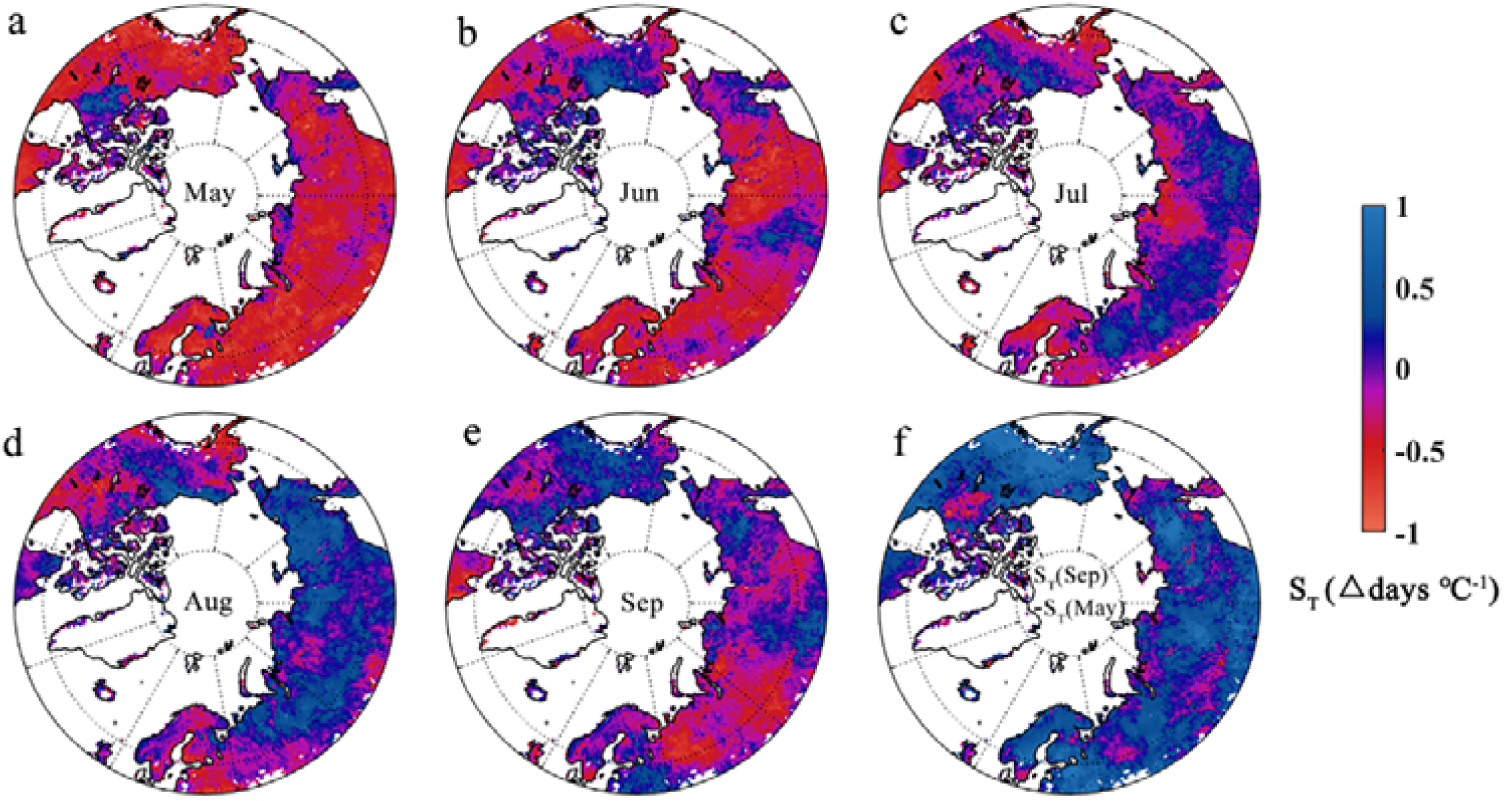
Spatial distribution of the monthly temperature sensitivities (S_T_, change in days □^−1^) of leaf senescence between May and September in the Northern Hemisphere. The calculated S_T_ values were based on phenological metrics extracted from the MODIS phenology product (MCD12Q2 version 6). (**a-e**), monthly S_T_ from May to September, **(f)** difference in the S_T_ between May and September.

**Fig. S5.**
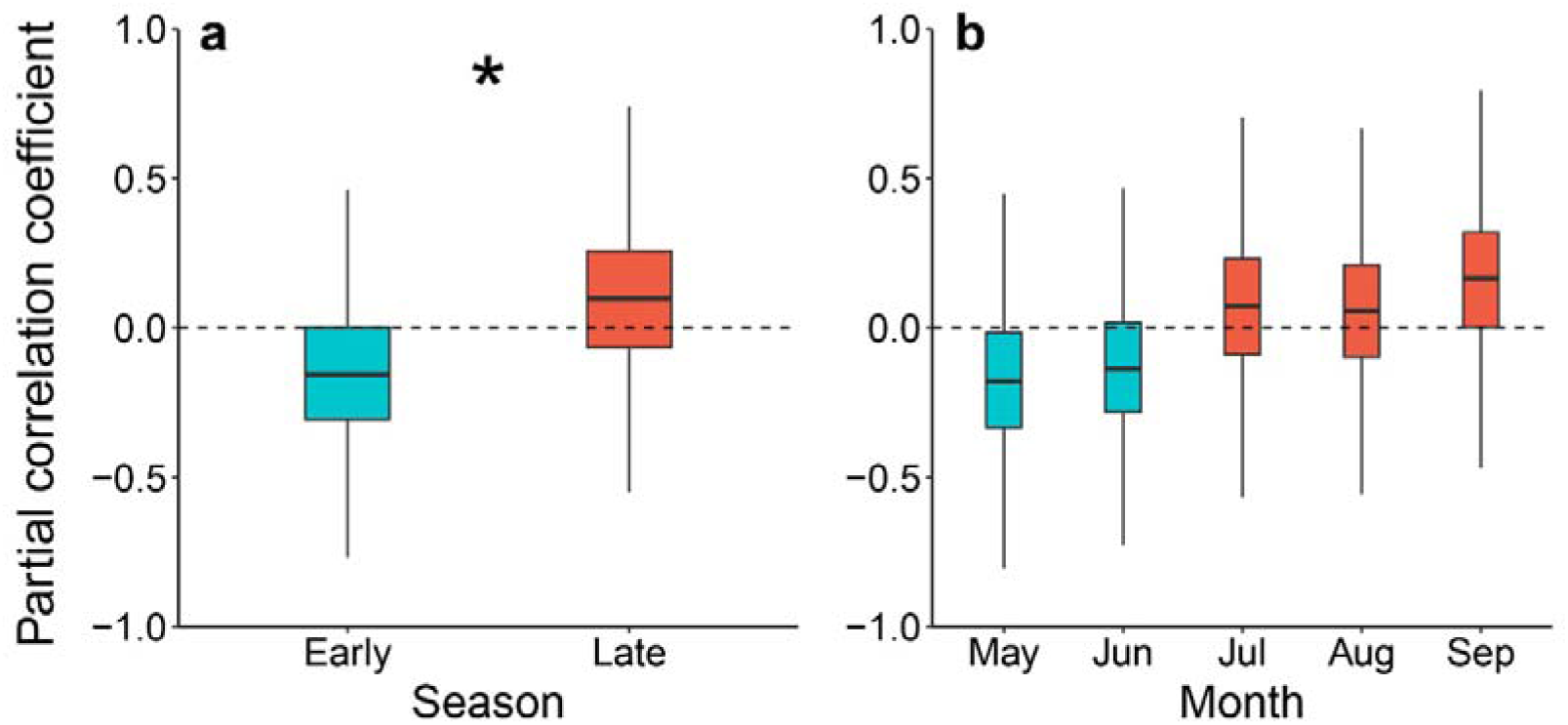
Partial correlation coefficients between temperature and leaf senescence dates during the early (May-June) and late season (July-September). The length of each box indicates the interquartile range, the horizontal line inside each box the median, and the bottom and top of the box the first and third quartiles respectively. The asterisk in **(a)** indicates a significant difference between the early and late season (*P*<0.05).

**Fig. S6.**
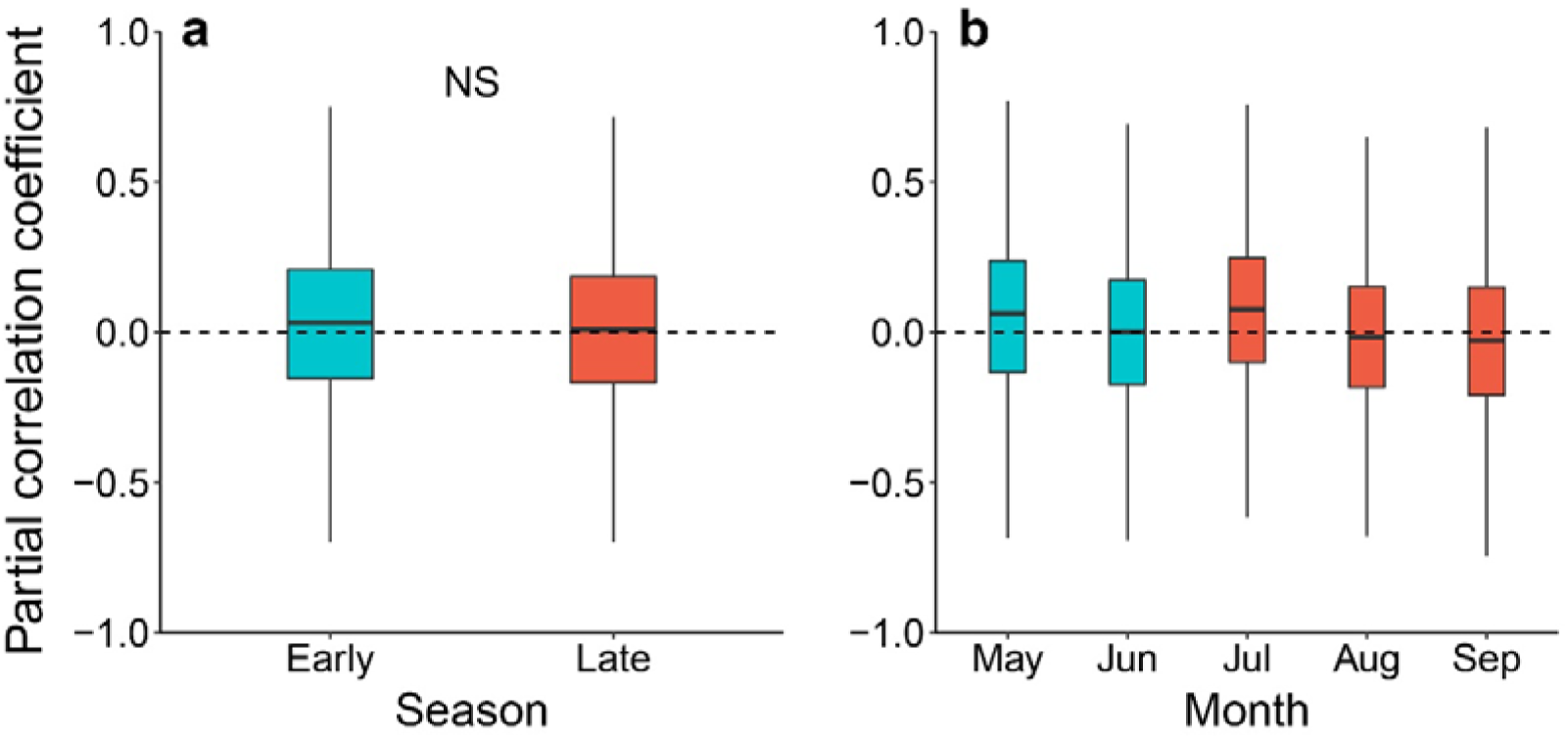
Partial correlation coefficients between soil moisture and leaf senescence dates during the early (May-June) and late season (July-September). The length of each box indicates the interquartile range, the horizontal line inside each box the median, and the bottom and top of the box the first and third quartiles respectively. The “NS” in **(a)** indicates no significant difference between the early and late season (*P*<0.05).

**Fig. S7.**
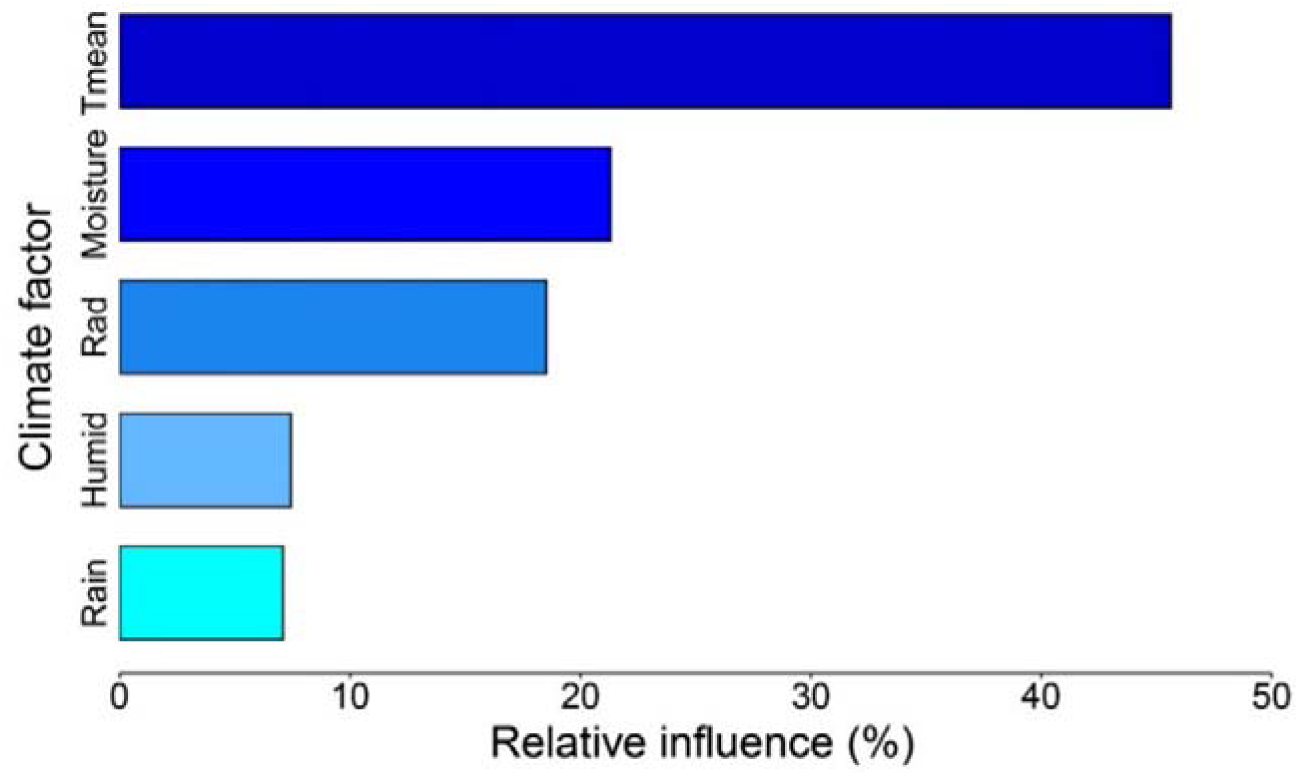
Relative influences of climate variables on leaf senescence dates during the growing season. The climate variables include mean temperature, soil moisture, radiation, humidity and precipitation between May and September.

**Fig. S8.**
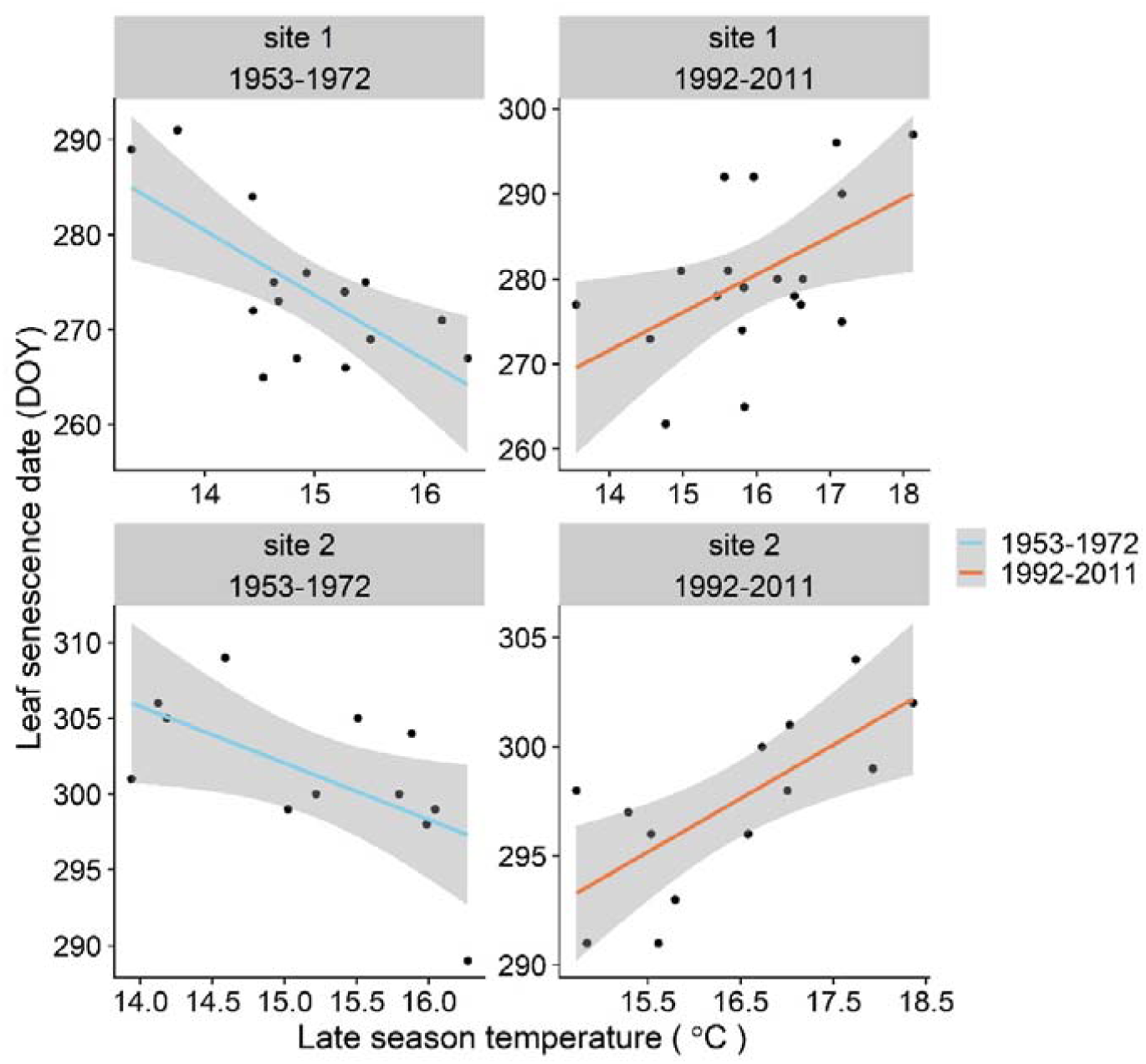
Effects of mean temperature during the late season (July-September) on leaf senescence dates of *Fagus sylvatica* (European beech) during 1953-1972 and 1992-2011 at several sites selected from the Pan European Phenology (PEP725) network. The leaf senescence date was expressed as the day of year (DOY). The shaded area indicates the 95% confidence intervals of the fitted regression lines.

